# The genetic origin of Daunians and the Pan-Mediterranean southern Italian Iron Age context

**DOI:** 10.1101/2021.07.30.454498

**Authors:** Serena Aneli, Tina Saupe, Francesco Montinaro, Anu Solnik, Ludovica Molinaro, Cinzia Scaggion, Nicola Carrara, Alessandro Raveane, Toomas Kivisild, Mait Metspalu, Christiana L Scheib, Luca Pagani

## Abstract

The geographical location and shape of Apulia, a narrow land stretching out in the sea at the South of Italy, made this region a Mediterranean crossroads connecting Western Europe and the Balkans. Such movements culminated at the beginning of the Iron Age with the Iapygian civilization which consisted of three cultures: Peucetians, Messapians and Daunians. Among them, the Daunians left a peculiar cultural heritage, with one-of-a-kind stelae and pottery, but, despite the extensive archaeological literature, their origin has been lost to time. In order to shed light on this and to provide a genetic picture of Iron Age Southern Italy, we collected and sequenced human remains from three archaeological sites geographically located in Northern Apulia (the area historically inhabited by Daunians) and radiocarbon dated between 1157 and 275 calBCE. We find that Iron Age Apulian samples are still distant from the genetic variability of modern-day Apulians, they show a remarkable genetic heterogeneity, even though a few kilometers and centuries separate them, and they are well inserted into the Iron Age Pan-Mediterranean genetic landscape. Our study provides for the first time a window on the genetic make-up of pre-imperial Southern Italy, whose increasing connectivity within the Mediterranean landscape, would have contributed to laying the foundation for modern genetic variability. In this light, the genetic profile of Daunians may be compatible with an autochthonous origin, with plausible contributions from the Balkan peninsula.

## Introduction

The Mediterranean’s Iron Age populations, between 1100 and 600 BCE, lived in a time of previously unprecedented connectivity^1^. Although the technological advances in seafaring had allowed great opportunities for long-distance mobility just a few millennia earlier, it was during the Iron Age that the cosmopolitan role of Mare Nostrum arose, by promoting the spreading of cultures, goods, languages, technical advances as well as heterogeneous ancestral genetic components coming from far and wide^1, 2^. Notable examples of such distant connections are the Greek and Phoenician settlements across the Central and Western Mediterranean shores beginning from the ninth and eighth centuries^3, 4^.

The Italian peninsula and its Tyrrenian islands, where the Iron Age is conventionally marked as beginning 950 BCE^1, 5^, joined as well the cosmopolitan wave sweeping across Southern Europe, by hosting numerous trading posts along its shores. A patchwork of communities appeared in this period within the Italian borders, each characterized by unique and well-defined cultures and identities, which were later encapsulated and blurred by the Roman colonization. The shift from Republican to Imperial Rome, with the consequent inclusion of non-Roman civilizations from and beyond the Italian peninsula, has already been shown to be connected with a major eastward genetic shift in Central Italian samples^6^. Genetic material from Imperial Romans from Central Italy is indeed the first to co-localize with contemporary Italians in a Principal Component space, pointing to this time period (∼200 BCE) as a crucial one in shaping contemporary Italian genetic makeup. Whether such a shift can also be observed in the rest of the Italian peninsula and how the Republic to Imperial transition may have impacted on the genetic landscape of local, pre-Roman populations remains an open question.

Despite the numerous written records and archaeological findings, questions about Iron Age populations, their origins and mutual relationships remain. Among the many groups occupying Italy in the Iron Age, the Daunians, a Iapygian population from northern Apulia, were first mentioned in the 7th-6th century BCE^7, 8^. Similarly to their neighbouring populations, Peucetians and Messapians (living in central and southern Apulia, respectively), the name of the Daunians comes from ancient Greek documents and, given the absence of written Daunian records, the scant information we have on their social, political and religious life are wholly reliant upon the material record, such as their one-of-a-kind stelae^8^. For instance, we know that they were mainly farmers, animal breeders, horsemen and maritime traders with an established trade network extending across the sea with Illyrian tribes^8–10^. A fascinating aspect of this population, as opposed to their neighbours in Apulia, was their tenacious resistance to external influences. For instance, they did not acquire either social or cultural Hellenic elements and no Greek alphabet inscriptions have been found in their settlements. Indeed, they retained a strong cultural identity and political autonomy until the Roman arrival in the late 4th - early 3rd century BCE^8^.

Despite the extensive archaeological literature, their origin has been lost to time and, as early as in the Hellenistic period, various legends already existed connecting them to either Illyria (an ancient region broadly identifiable with the Balkan peninsula), Arkadia (present-day Peloponnesus) or Crete^7, 8^.

To shed light on this population and provide a glance of the Iron Age genetic landscape of the Southern Italian peninsula, we collected human remains from three Iron Age necropoleis geographically located within the area historically inhabited by Daunians: *Ordona* (ancient Herdonia), one of the largest settlements estimated at 600 hectares^8^, *Salapia,* and *San Giovanni Rotondo* (Figure 1A). Here, we provide for the first time a genetic investigation of Iron Age Apulian associated with the Daunian culture, offering insight into the genetic landscape of pre-Imperial Southern Italy.

**Figure 1:**
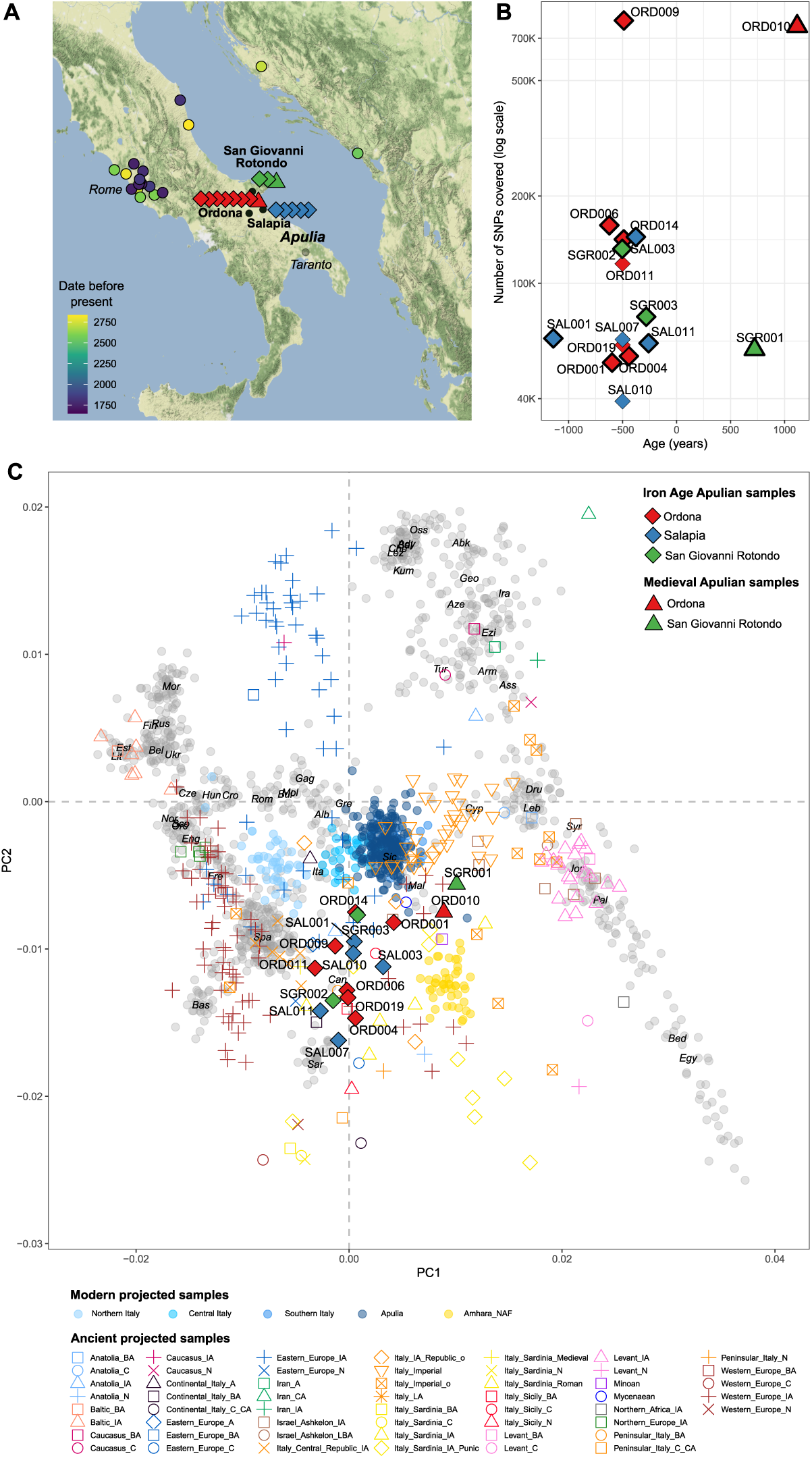
Geographical location, dating and genetic structure. (A) Geographical location of the newly generated Apulian ancient individuals (green, red and blue diamonds and triangles) and other published Iron Age samples from Italy and surrounding areas (color scale mirroring their “years before present” dates). See also Table 1, Data S1, and Data S2. (B) Dating and number of SNPs from 1240K set covered. Samples with thick edges have been radiocarbon dated (mean presented), while for the others we represented their archaeological dating (Table 1, Data S1B). (C) Principal component analysis (PCA) of newly generated individuals with reference ancient individuals projected onto the genetic variation inferred from modern-day Western Eurasian populations (Data S2 and Data S3). For the ancient reference populations which have not been attributed to the Iron Age, we plotted their centroids.

**Table 1.**
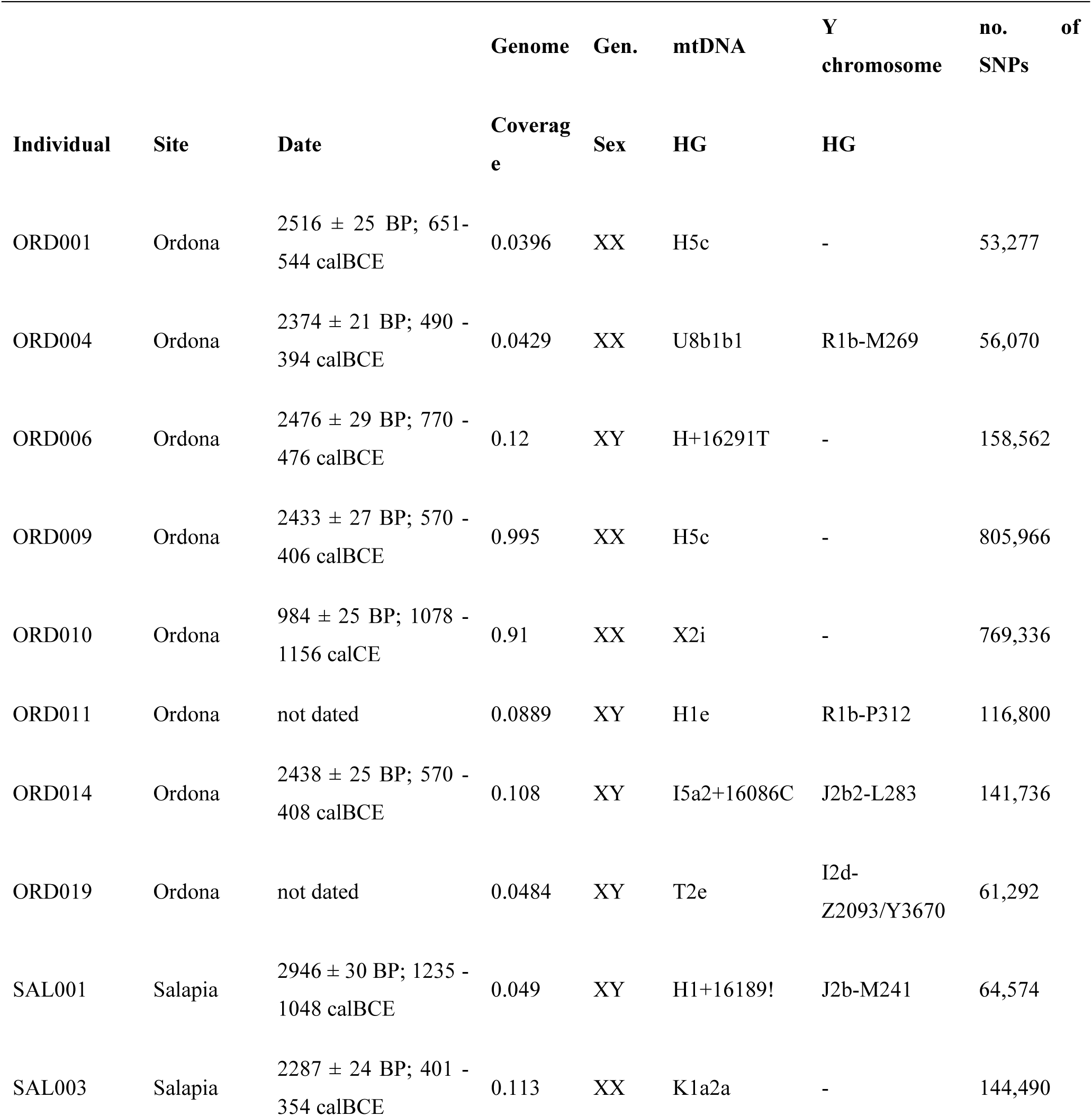
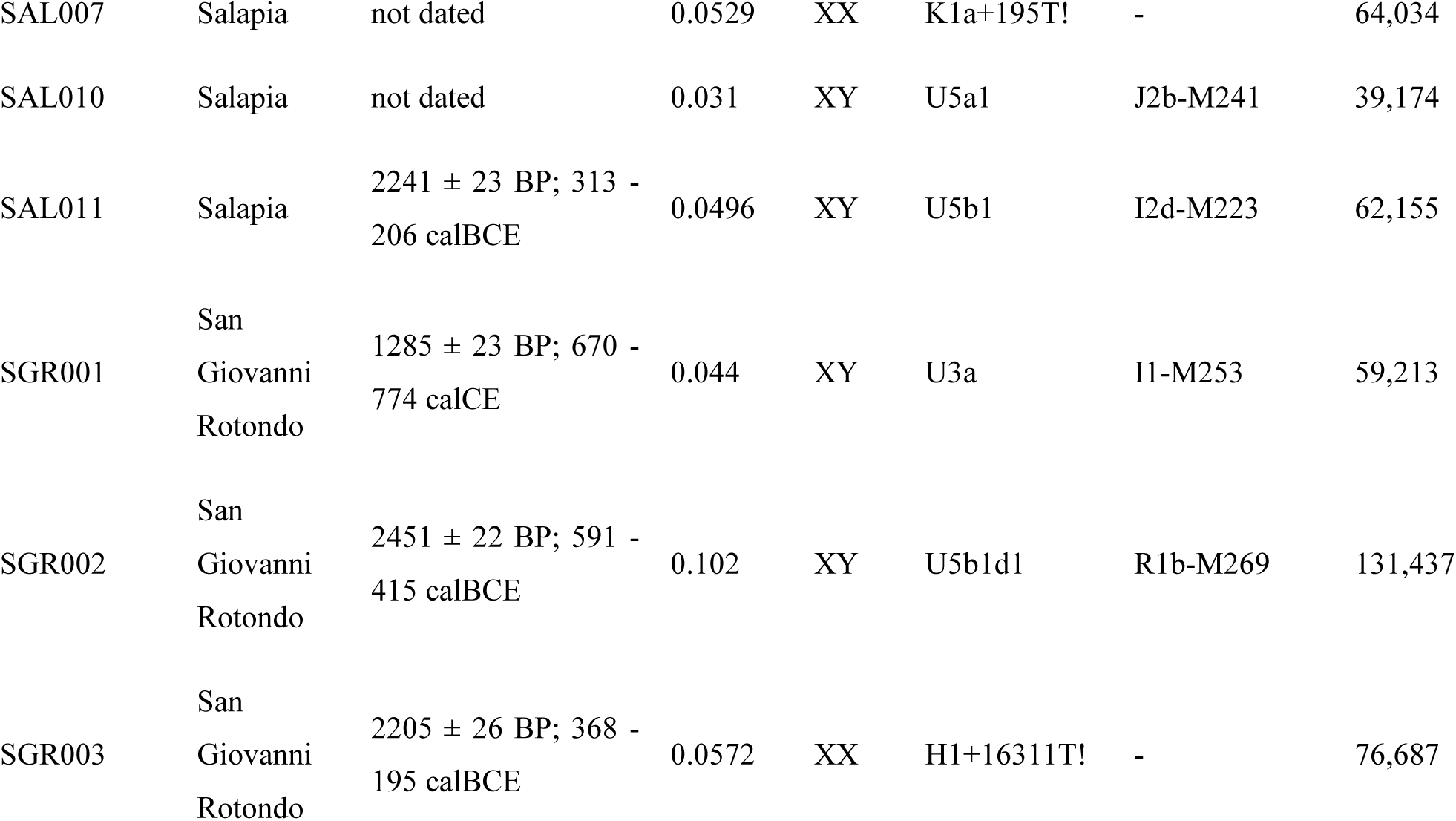
Archaeological information, dating, genome coverage, genetic sex, mtDNA, and Y chromosome haplogroups of the individuals of this study selected for genome-wide analysis. Dates in BP are raw radiocarbon dates, calibrated dates are the 95.4% probability and were calibrated using IntCal20^30^. Gen.-genetic; HG-haplogroup. See also Data S1.

## Results

We extracted DNA from 34 human skeletal remains (petrous bone = 23 and teeth = 11) from three necropoleis (Ordona = 19; Salapia = 12; San Giovanni Rotondo = 3; hereafter ORD, SAL and SGR, respectively) at the Ancient DNA Laboratory of the Institute of Genomics, University of Tartu in Estonia (Data S1A). All three necropoleis are geographically located less than 50 km from each other in modern Apulia, Southeastern Italy. Ordona and Salapia are archaeologically dated to the Daunian period (6th-3rd century BCE) except one individual (ORD010) archaeologically dated to the Medieval time period (which in Europe dates approximately from the 5th to the late 15th century CE) (Figure 1B). Based on the museum record, the samples from the San Giovanni Rotondo necropole have been archaeological inferred to be from the Iron Age period. After screening 21 libraries at a low depth (±20M reads per library), we additionally sequenced to higher depth 16 libraries with endogenous DNA between 1.81 - 38.82% and mitochondrial DNA-based (mtDNA) contamination estimated at less than 2.89% (Data S1A).

The sequencing runs were merged resulting in 16 individuals for genome-wide analysis: eight from Ordona, five from Salapia, and three from San Giovanni Rotondo. The final dataset includes individuals with an average genomic coverage between 0.031 - 0.995× and a number of single nucleotide polymorphisms (SNPs) overlapping with Human Origins 1240k between ∼40,000 and ∼810,000 (Data S1A).

Out of those 16 individuals, we selected 10 individuals based on their proximity within the principal component analysis (PCA) space (Figure 1C) for radiocarbon dating and estimated their age between 1157 and 275 calBCE with a median date of 521 calBCE (Data S1B). The radiocarbon dates confirm the archaeological dates of individuals from Salapia and Ordona as well as the Iron Age affiliation of the two San Giovanni Rotondo samples (SGR002 and SGR003). Two additional samples (ORD010 and SGR001) with a shift towards the Near East in the PCA (Figure 1C) were radiocarbon dated to 1078 - 1156 calCE (95.4%) and 670 - 774 calCE (95.4%) respectively (Data S1B). Based on their dates, the samples were used as an external control for further analyses focusing on Iron Age Apulia (IAA).

We determined the mitochondrial DNA (mtDNA) haplotype of each individual and the Y chromosome (Ychr) haplogroup for 9 male individuals (See STAR methods and Data S1A and S1C-D). We found that the mtDNA haplotypes mostly belong to the mtDNA lineages H1, H5, K1, and U5, haplotypes found in previous studies of individuals from this time period in the Italian Peninsula^6^. Besides the Ychr haplogroup R1b, which is the most frequent haplogroup during the Bronze Age in the Italian Peninsula and on the islands Sardinia and Sicily^4, 6, 11–13^, we found the Ychr lineages I1-M253, I2d-M223, and J2b-M241. The haplogroup I2d-M223 was one of the main Y chromosome lineages in Western Europe until the Late Neolithic whereas J2b-M241 first appears in the Bronze Age^3, 4, 12, 14, 15^. We found one Early Medieval individual (SGR001 (670 - 774 calCE (95.4%)) belonging to haplogroup I1-M253, which is common in Northern Europe and previously also detected in a 6th Century Langobard burial from North Italy ^16^.

We used READ^17^ to identify pairs with first or second degree of genetic relatedness from autosomal data (See STAR methods and Data S1E-H for details, Figure S11). We found one first-degree relationship between the two female individuals ORD001 (age: 9 - 11 years, 651- 544 calBCE (95.4%)) and ORD009 (age: 40 - 45 years, 570 - 406 calBCE (95.4%)) from Ordona sharing the identical mtDNA haplogroup H5c and indicating a mother-daughter relationship (Data S1, Figure S1A-B).

### The making of modern Italians

To explore the genetic make-up of the IAA population, we performed a PCA projecting the ancient individuals onto the genetic variation of modern Eurasian samples (Figure 1C, Data S2 and Data S3). Our samples are largely scattered between modern peninsular Italians and Sardinians, and, in contrast to what was generally described^18, 19^ for other European Iron Age populations (e.g., Northern_Europe_IA, Western_Europe_IA and Levant_IA in Figure 1C), they are still clearly distant from the genetic variability of modern-day inhabitants of Apulia. The downward shift of Iron Age Apulians from the present-day ones is further confirmed by the significantly negative *f*4*(Modern Apulians, IAA; X, Mbuti)*, where *X* is a Neolithic/Chalcolithic/Copper Age population (Figure 2A, Data S4). Within the remarkable heterogeneity reported by the PCA, which does not mirror the archaeological sites, the two medieval individuals are shifted towards modern Middle Eastern and Caucasus populations (ORD010 and SGR001), while the others are stretched along the PC2. This pattern partially mirrors the chronological date with the most recent being more similar to present-day Southern Europeans, and is further strengthened when considering the PC3 distribution (Figure S2). Three samples located at the bottom of the PCA (ORD004, ORD019, SAL007) and one (SAL010) falling in the middle did not include modern Apulians among the top 25 results of an *f*3 outgroup analysis (Figure S3). All of them showed an affinity to Copper and Bronze Age Italians^11^ as well as the Aegean and the Mediterranean worlds (including Minoans, Greece, Croatians, and Gibraltar). A similar distribution is mirrored in the Multi-Dimensional scaling (MDS) built from the *f*3 outgroup measures, where the oldest IAA individual (SAL001; 1235 - 1048 calBCE (95.4%)) lies farthest from the modern samples, while the medieval ones (ORD010: 1078 - 1156 cal CE (95.4%) and SGR001: 670 - 774 cal CE (95.4%)) are the closest (Figure S4).

**Figure 2:**
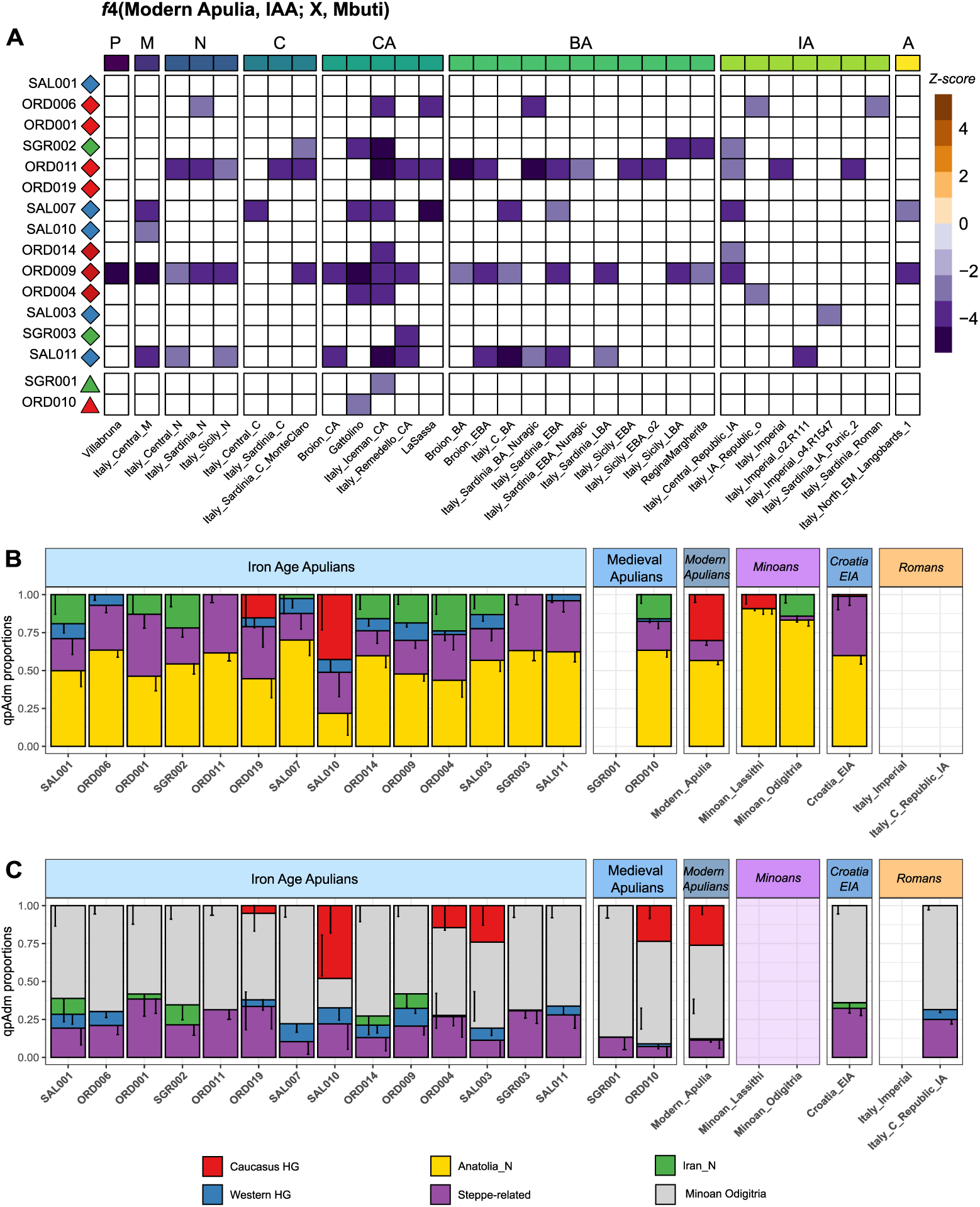
Genetic relationship of Iron Age and Middle Age Apulians and their ancestral composition. (A) Heatmap representing the Z-score values of f4(Modern Apulia, IAA; X, Mbuti) where X is an ancient Italian population. We also added the two Middle Age samples for comparison. Tests with Z-scores between -3 and 3 or with less than 5,000 SNPs were not included (P: Palaeolithic, M: Mesolithic, N: Neolithic, C: Chalcolithic, CA: Copper Age, BA: Bronze Age, IA: Iron Age, A: Antiquity). (B) qpAdm proportions of ancient Apulian samples and other reference populations (Modern Apulians, Minoans, Croatia EIA and the Roman individuals from the Republican and the Imperial period) using “base” sources: Caucasus and Western hunter-gatherer component (HG), Anatolia Neolithic (Anatolia N), Iranian Neolithic (Iran N), Stepperelated ancestry. (C) qpAdm proportions using also the Minoans as source (STAR methods). In B and C, we plotted the model with the highest number of sources and p-value (complete results may be seen in Figure S5).

The peculiar positioning of the IAA individuals casts doubt on when the major population shift resulting in modern Italian genetic composition took place. The shift towards the modern Italian genetic variability can be seen with the Republican-Imperial Roman samples^6^, the latter being more “similar” to modern Italians (Figure 1C). Whether Apulian individuals dating back to the Imperial phase would also show a repositioning towards modern genetic variability remains an open question, although the later, medieval samples of this study point in that direction.

### The pan-Mediterranean genetic landscape of Iron Age Apulia

The geographic location of Apulia, a narrow peninsula stretching out in the sea at the South of Italy, has made this region an important Mediterranean crossroads connecting Western Europe, the Balkans, the Aegean, and Levant worlds. This is reflected in the PCA where IAA individuals are closely related to other Iron Age populations from the Mediterranean and surrounding areas (e.g., Montenegro, Bulgaria and Sardinia) (Figure 1C and Figure S4). Nomadic or cosmopolitan groups scatter like IAA: three Punic individuals from Sardinia (Italy_Sardinia_IA_Punic^3^), three Moldova Scythians already reported to be genetically similar to Southern Europeans^20^, Spanish individuals from the Hellenistic and the Romans periods^21^ and an individual from the 12^th^ century Iron Age Ashkelon^22^ which clusters with ORD001.

In order to shed light onto the genetic composition of the IAA individuals, we modelled them as a combination of the main ancestries documented across Western Europe at that time: Western Hunter-Gatherers (WHG), Anatolian Neolithic (AN), Steppe-related and, interchangeably, Caucasus Hunter-Gatherers (CHG) or Iranian Neolithic (IN) using the qpWave/qpAdm framework (Figure 2B, Figure S5A, STAR Methods). Broadly, the contributions of such ancestries to the genetic variability of ancient European populations vary according to their geographical positions: in particular, northernmost locations received higher proportions of WHG, Steppe-related ancestry and, consequently, CHG ancestries, while Southern European groups carried variable Iranian Neolithic or CHG traces^23^. In view of this, we observed that while the IAA individuals could generally be modelled as a two-way admixture between AN and Steppe (0.63±0.08 and 0.37±0.08, respectively), the alternative model AN + CHG/IN could also fit for a subset of them, particularly in case of the samples ORD004, ORD010 and SAL010 with higher or comparable p-values (Figure S5A, first row with two sources). When three or four sources were tested, the presence of WHG ancestry in the majority of our individuals emerges, which, together with AN, Steppe and CHG/IN, forms a supported model for IIA samples (Figure 2B, Figure S5A and Data S5). Notably, for the individuals stretching downwards in the PCA (ORD004, ORD019, SGR002 and the Medieval ORD010) a three-way admixture involving AN, Steppe and CHG/IN is generally preferable. To better understand the putative contribution of more recent populations, we modelled our samples with base sources (WHG, AN, Steppe-related and CHG/IN) and, alternatively, Minoans (Odigitria and Lasithi), Amhara_NAF and Roman Republicans (Figure 2C, Figure S5B-D, STAR Methods). Amhara_NAF can be used as a proxy for the Non-African component in modern Ethiopian individuals that was tentatively linked to the Sea People, a Bronze Age nomadic seafaring population^22, 24^. Together with Minoans and Roman Republicans, this component can be broadly modelled as a Pan-Mediterranean population (constituted by AN and IN/CHG components) with the addition of WHG and Steppe-related ancestry in Roman Republicans. When modelled also with Minoans and Amhara_NAF, which roughly proxies the same ancestral signature, the majority of the samples required an additional CHG/IN contribution (two-way admixtures in Figure S5B,C) as well as Steppe-related and WHG. We further observed that, as previously seen, the WHG contribution is less clear in those samples stretching downwards in the PCA. While the CHG/IN additional contribution may simply proxy the presence of Steppe-related ancestry in IAA, the absence of which in Minoans has already been reported^25^, the same can not be said about Roman Republicans (two-way admixtures in Figure S5D), which harboured a considerable amount of Steppe component^6^. However, this signature is not confirmed with *f*4 analyses (Figure S6 B,C), where just Mycenaean groups report less CHG ancestry than our samples.

The broader picture emerging from qpAdm analyses is the pervasive presence of coeval Italian ancestries (Rome Republicans), as well as previous Bronze Age sources (Minoans and Sea People) spread far and wide across the Mediterranean Sea. In such a melting pot scenario, the peculiar genetic heterogeneity of IAA individuals, who lived in close temporal and geographical proximity, stand out (Figure 1C, Figure S4, Figure S7). An example of the cosmopolitan nature of Iron Age Mediterranean is the first-degree relationship between ORD009 (mother) and ORD001 (daughter), whose positions in the PCA strikingly differ with the individual ORD001 being stretched towards Middle Eastern and Caucasus modern populations, as a consequence of the foreign origin of the father (Figure 1C and Figure S4). *F*4 analyses in the form *f4(ORD009, ORD001; X, Mbuti)* report a slightly significant (Z-score between -2 and -3) excess of Greece_N, Portugal_LN_C, Lebanon_Roman and Italy_Sicily_EBA in ORD001, which may explain its eastward shift (Figure S8A). Moreover, ORD001, together with SGR002, but not ORD009 harboured more CHG when compared to Lebanon_Hellenistic and Lebanon_IA3 samples, respectively (Figure S8C), as well as an increase in Lebanon_Hellenistic traces when compared with modern Apulians (data not shown).

We also investigated whether the PCA scattering was due to varying African or Levantine contributions with *f*4(*Rome Republican, IAA, Levant_N/YRI, Mbuti)* and tried the same on Medieval ancient Apulians (ORD010 and SGR001). However, none of the tested ancient Apulians shows a significant excess of YRI ancestry when compared to the contemporary Roman Republicans, even though ORD014, SAL007 and SAL011 show negative *f*4 values with a Z-score between 2 and 3 (Figure S8B).

### The origin of Daunians

The genetic heterogeneity of IAA individuals connected to the Daunian population and the cosmopolitan landscape of Iron Age Mediterranean populations hinder a full reconstruction of the demographic processes leading to the Daunians. Nevertheless, a few milestones can be spotted.

When we performed *f*3 analyses to investigate the nearest possible source for each IAA individual using Minoans, Iron Age Croatians and the local Roman Republicans (Figure 3A), we found that none of the IAA individuals shows higher affinities with Minoans. Three of them, clustered close to modern Italians in the PCA (ORD001, ORD014 and SGR003, Figure 1C), show higher affinity with the Iron Age Croatian sample (ORD004 followed this pattern too, but with lower *f*3 values). However, the remaining majority are closest to the Roman Republicans, which can be interpreted as representative of local Iron Age peninsular Italy ancestry.

**Figure 3:**
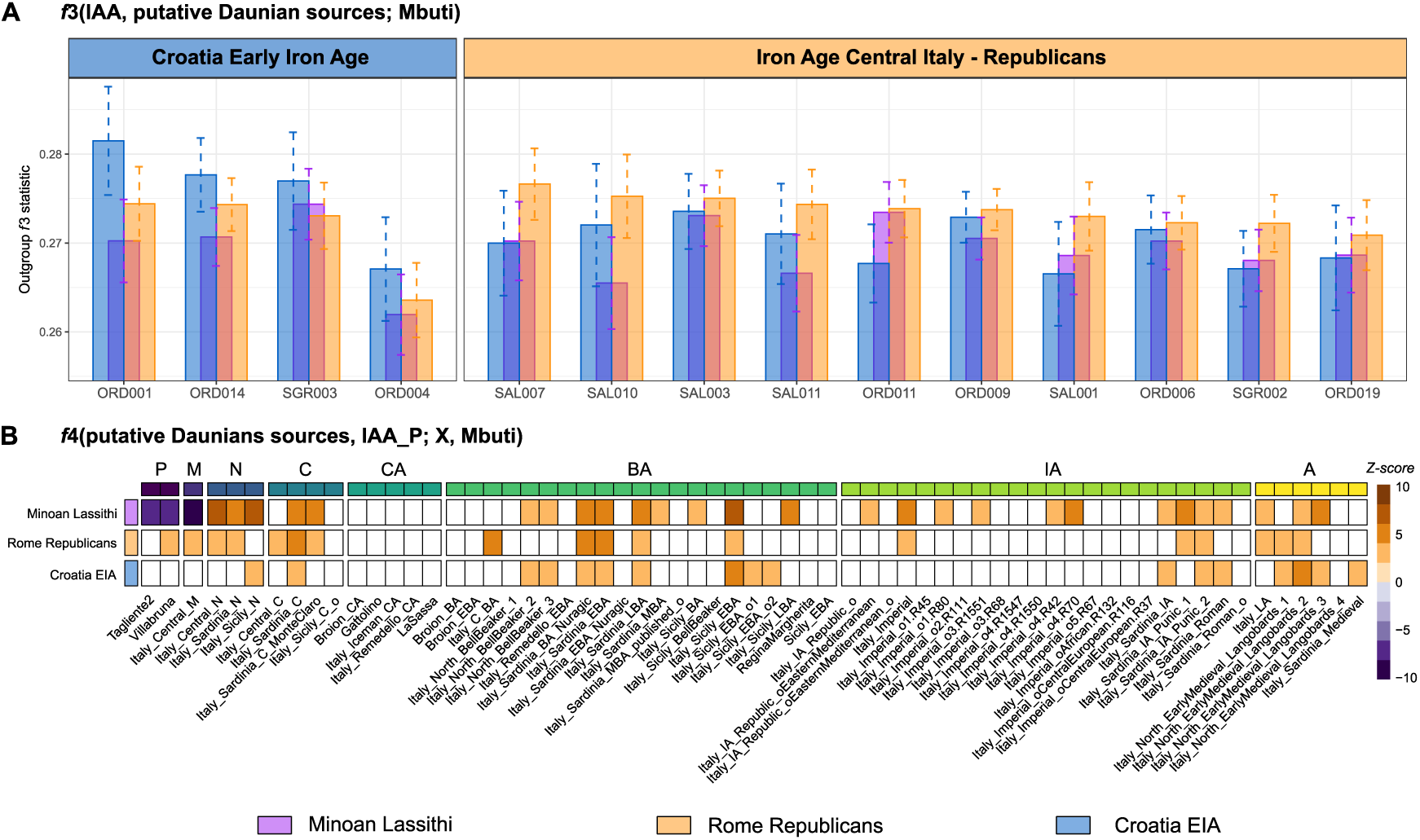
Genetic affinities of Iron Age Apulian samples with the putative populations of origin: Minoans (Minoan Lassithi), Illyrians (here proxied by the Croatia EIA individual) and the Roman Republicans (here proxying the autochthonous Iron Age Italian ancestry). (A) Outgroup f3 statistics of IAA samples compared with the putative Daunian sources. Samples have been sorted and grouped according to the source whose f3 values were higher. (B) Heatmap showing the Z-score values of f4(putative sources, IAA P; X, Mbuti) where X is an ancient Italian population and IAA P is the entire set of Iron Age Apulian samples taken together. Tests with Z-scores between -3 and 3 or with less than 5,000 SNPs were not included (P: Palaeolithic, M: Mesolithic, N: Neolithic, C: Chalcolithic, CA: Copper Age, BA: Bronze Age, IA: Iron Age, A: Antiquity).

Moreover, the WHG contribution, which was a necessary component to explain IAA and Roman Republicans according to qpAdm output, is absent from Minoans and Iron Age Croatians, thus making it a putative signature of an at least partial autochthonous origin (Figure 2B,C and Figure S5). These results are confirmed for Minoans, but not for Croatia_EIA, by the *f*4 analyses (Figure 3B, Figure S6A and Data S6). Indeed, the significantly negative *f*4(*Minoans, IAA; WHG, Mbuti)* (Figure 3B and Figure S6A), as well as the *f*4(*Greece_Minoan_Lassithi/Greece_BA_Mycenaean/Greece_BA_Mycenaean_Pylos, IAA; X, Mbuti),* reported a significant excess of WHG ancestry (*X* being Tagliente2, Loschbour, Italy_Central_M, Switzerland_Bichon and so on) in all IAA individuals, with the exception of ORD004, SAL001 and SGR003 (also the medieval samples ORD010 and SGR001 deviate from this pattern, Data S6). An excess of Bronze Age Steppe-related component, which at that time was already present along the Italian peninsula^11^, was clearly present in SGR002, ORD006 and ORD009 by the *f*4(*IAA, Greece_Minoan_Lassithi; X, Mbuti*) (Data S6).

The excess of WHG ancestry, tentatively suggesting a local origin, is somewhat blurred by the genetic similarity of the two most probable sources - Illyrians (Croatia EIA) and an autochthonous one (Roman Republicans), which together make part of the same Mediterranean continuum. Indeed, while no significantly negative *f*4*(Croatia_EIA, IAA; X, Mbuti)* values have been found (with the exception of *X* being Italy_C_BA, Italy_Iceman_CA, Portugal_MBA and Anatolia_MLBA in SAL007, which retains the highest proportion of AN in qpAdm (Figure 2B and Figure S5), the same pattern is obtained when IAA individuals are compared with Roman Republicans (*f*4*(Roman Republicans, IAA; X, Mbuti)*, (Data S6)). Imperial individuals from a few centuries later (*f*4(*Rome_Imperial, IAA; X, Mbuti*)) show quite the opposite pattern: no positive results have been produced with the exception of ORD004 (X=Lebanon_Roman), SAL001 (X=Italy_Sardinia_Roman_o), SAL010 (X=Russia_LateMaikop) and ORD010. Conversely, the negative *f*4 values point toward WHG, Neolithic and Bronze Age Steppe-related ancestries (Data S6).

Another signal coming from qpAdm analyses is the apparent excess of CHG ancestry in IAA; however, the predominant contribution of CHG to the Steppe-related ancestry that, by the Iron Age, had already spread to the Mediterranean area makes it hard to properly detect a CHG signature independent from the Steppe wave, possibly brought by pan-Mediterranean influxes. When directly investigated with an *f*4 framework, IAA shows generally more CHG than Mycenaean, less CHG than contemporary Croatian_EIA and, in some cases (ORD019, SGR002 and the Medieval SGR001 with Z-scores higher than 2) more CHG than older Croatian samples (_N and _MN) (Figure S6).

## Discussion

The new genomic sequences Daunian samples reveal that Iron Age (pre-Imperial) Southern Italy (Apulia) can be placed within a Pan-Mediterranean genetic continuum that stretches from Crete (Minoans^25^) and the Levant (Sea People^22, 24^) to the Republican Rome and the Iberian Peninsula^6^, mainly composed by AN and IN/CHG genetic features with the addition of WHG and Steppe-related influences in Continental Italy. Pre-Imperial Italian populations, being part of this broader landscape, are not directly superimposable with contemporary Italians which instead seem to be influenced by the homogenizing effect of Imperial Roman and late Antiquity events.

Within the described Pan Mediterranean landscape, the IAA/Daunians show a compelling heterogeneity, and the highest genetic affinity to Republican Romans and Iron Age Croatians, while Minoans and other Iron Age Greek samples show absent or reduced WHG contribution when compared to IAA. This makes a Cretan or Arkadian origin less likely, even though some tales have connected them with the Greek hero Diomedes and many ancient historians have claimed such origins for their neighbour Messapians and Peucetians.

The Daunians maintained strong commercial and political relations with the Illyrian people, controlling together the area spanning from the Dalmatia to the Gargano peninsula^10^ and had many cultural affinities with them^26^. The material culture, involving peculiar anthropomorphic statue stelae, has provided some information on Daunian culture and may also help in unravelling their mysterious origin. In particular, the forearm decorations on a female stela have been interpreted as tattoos and, while tattooing practices were considered barbarian among the Greeks^27^, they were customary in populations from Tracia and Illyria and, more generally, among the women of status from the Balkans^8, 28^.

It is not clear whether these connections indicate a movement of people or a sharing of cultural ideas and a conclusive answer to the origin of the Daunians remains elusive. From a parsimony perspective, the genetic results point to an autochthonous origin, although we cannot exclude additional influences from Croatia (ancient Illyria), as described by available historical sources and by the material remains^8, 29^.

## STAR Methods

### Resource availability

#### Lead Contact

Further information and requests for resources and reagents should be directed to and will be fulfilled by the Lead Contact: Serena Aneli

### Materials Availability

This study did not generate new unique reagents.

### Data and Code Availability

The DNA sequences generated during this study are available at the European Nucleotide Archive (ENA; [link]) at the accession number [accession number]. The data are also available in PLINK format through the data depository of the EBC (http://evolbio.ut.ee).

### Experimental model and subject details

The ancient human remains analysed in this work belong to the collections of the Museum of Anthropology, University of Padua and have been found and catalogued by different archaeological campaigns carried out during the twentieth century.

### Ordona

The necropolis of Herdonia (today’s Ordona, within the Apulian Foggia province in Italy) was studied by different archaeological campaigns interested in the Daunians, Roman as well as the medieval settlements in 1978 and 1981^31^. The human remains collected on such occasions were later extensively catalogued and studied and new paleopathological evidence was also brought back to light^32^. Inhabited starting from the Neolithic period, Herdonia became an important Daunian center from the 6th century BCE.

For this study, in collaboration with the University of Padova and Museum of Anthropology of Padova, samples of human remains (petrous bone=19) were taken. The samples ORD001, ORD004, ORD006, ORD009, ORD010, and ORD014 were dated at ^14^CHRONO Centre for Climate, the Environment, and Chronology in Belfast, UK (Data S1B).

### Salapia

The necropolis of Salapia, an ancient town located 10 km from contemporary Cerignola, within the Apulian Foggia province in Italy. The osteological samples were brought by Prof. Santo Tiné to the Museum of Anthropology of the University of Padova with no further information on the archaeological context, and osteological studies were carried out by Cleto Corrain and colleagues in 1971^33^. From this site, samples of 12 human remains (petrous bone=5, teeth=7) were taken. The samples SAL001, SAL003, and SAL011 were dated at ^14^CHRONO Centre for Climate, the Environment, and Chronology in Belfast, UK (Data S1B).

### San Giovanni Rotondo

The San Giovanni Rotondo samples come from the osteo-archaeological collection of the Museum of Anthropology of the University of Padova and are not associated to any further record with the exception of a broad “Iron Age” archaeological label and may be part of the samples brought to the Museum by Prof. Santo Tiné in the 1960s. From this site, samples of human remains (petrous bone=3) were taken. The samples SGR001, SGR002, and SGR003 were dated at ^14^CHRONO Centre for Climate, the Environment, and Chronology in Belfast, UK (Data S1B).

### Method Details

All of the laboratory work was performed in dedicated ancient DNA laboratories at the Estonian Biocentre, Institute of Genomics, University of Tartu, Tartu, Estonia. The library quantification and sequencing were performed at the Core Facility of the Institute of Genomics, Tartu, Estonia. The main steps of the laboratory work are detailed below.

### DNA extraction

In total 34 samples from human remains were extracted for DNA analysis (Data S1).

The first layer of pars petrous was removed with a sterilised drill bite to avoid exogenous contamination. A 10 mm core of the inner ear was sampled from the pars petrous. The drill bits and core drill were sterilised in between samples with 6% (w/v) bleach followed by distilled water and then ethanol rinse. Root portions of teeth were removed with a sterile drill wheel.

The root and the petrous portions were soaked in 6% (w/v) bleach for 5 minutes. Samples were rinsed three times with 18.2 MΩcm H_2_O and soaked in 70% (v/v) Ethanol for 2 minutes. The tubes were shaken during the procedure to dislodge particles. The samples were transferred to a clean paper towel on a rack inside a class IIB hood with the UV light on and allowed to dry for two to three hours.

Afterwards, the samples were weighed to calculate the accurate volume of EDTA (20x EDTA [µl] of sample mass [mg]) and Proteinase K (0.5x Proteinase K [µl] of sample mass [mg]). EDTA and Proteinase K were added into PCR-clean 5 ml or 15 ml conical tubes (Eppendorf) along with the samples inside the IIB hood and the tubes were incubated 72 h on a slow shaker at room temperature.

The DNA extracts (of root portions and pars petrous portions) were concentrated to 250 µl using the Vivaspin® Turbo 15 (Sartorius) and purified in large volume columns (High Pure Viral Nucleic Acid Large Volume Kit, Roche) using 2.5 ml of PB buffer, 1 ml of PE buffer and 100 μl of EB buffer (MinElute PCR Purification Kit, QIAGEN). For the elution of the endogenous DNA, the silica columns were transferred to a collection tube to dry and followed in 1.5 ml DNA lo-bind tubes (Eppendorf) to elute. The samples were incubated with 100 µl EB buffer at 37 C for 10 minutes and centrifuged at 13,000 rpm for two minutes. After centrifugation, the silica columns were removed and the samples were stored at -20 C. Only one extraction was performed per extraction for screening and 30 µl used for libraries.

### Library preparation

Sequencing libraries were built using NEBNext® DNA Library Prep Master Mix Set for 454™ (E6070, New England Biolabs) and Illumina-specific adaptors^34^ following established protocols^34–36^. The end repair module was implemented using 18.75 μl of water, 7.5 μl of buffer and 3.75 μl of enzyme mix, incubating at 20 °C for 30 minutes. The samples were purified using 500 μl PB and 650 μl of PE buffer and eluted in 30 μl EB buffer (MinElute PCR Purification Kit, QIAGEN). The adaptor ligation module was implemented using 10 μl of buffer, 5 μl of T4 ligase and 5 μl of adaptor mix^34^, incubating at 20 °C for 15 minutes. The samples were purified as in the previous step and eluted in 30 μl of EB buffer (MinElute PCR Purification Kit, QIAGEN). The adaptor fill-in module was implemented using 13 μl of water, 5 μl of buffer and 2 μl of Bst DNA polymerase, incubating at 37 °C for 30 and at 80 °C for 20 minutes. Libraries were amplified using the following PCR set up: 50μl DNA library, 1X PCR buffer, 2.5mM MgCl2, 1 mg/ml BSA, 0.2μM inPE1.0, 0.2mM dNTP each, 0.1U/μl HGS Taq Diamond and 0.2μM indexing primer. Cycling conditions were: 5’ at 94C, followed by 18 cycles of 30 seconds each at 94C, 60C, and 68C, with a final extension of 7 minutes at 72C. The samples were purified and eluted in 35 μl of EB buffer (MinElute® PCR Purification Kit, QIAGEN). Three verification steps were implemented to make sure library preparation was successful and to measure the concentration of dsDNA/sequencing libraries - fluorometric quantitation (Qubit, Thermo Fisher Scientific), parallel capillary electrophoresis (Fragment Analyser, Agilent Technologies) and qPCR.

### DNA sequencing

DNA was sequenced using the Illumina NextSeq500/550 High-Output single-end 75 cycle kit. As a norm, 15 samples were sequenced together on one flow cell; additional data was generated for 16 samples to increase coverage (Data S1).

### Quantification and statistical analysis

#### Mapping

Before mapping, the sequences of the adapters, indexes, and poly-G tails occuring due to the specifics of the NextSeq 500 technology were cut from the ends of DNA sequences using cutadapt-1.11^37^. Sequences shorter than 30 bp were also removed with the same program to avoid random mapping of sequences from other species. The sequences were aligned to the reference sequence GRCh37 (hs37d5) using Burrows-Wheeler Aligner (BWA 0.7.12)^38^ and the command mem with re-seeding disabled.

After alignment, the sequences were converted to BAM format and only sequences that mapped to the human genome were kept with samtools 1.3^39^. Afterwards, the data from different flow cell lanes were merged and duplicates were removed using picard 2.12 (http://broadinstitute.github.io/picard/index.html). Indels were realigned using GATK 3.5^40^ and reads with a mapping quality less than 10 were filtered out using samtools 1.3^39^.

#### aDNA authentication

As a result of degradation over time, aDNA can be distinguished from modern DNA by certain characteristics: short fragments and a high frequency of C=>T substitutions at the 5’ ends of sequences due to cytosine deamination. The program mapDamage2.0^41^ was used to estimate the frequency of 5’ C=>T transitions. Rates of contamination were estimated on mitochondrial DNA by calculating the percentage of non-consensus bases at haplogroup-defining positions as detailed in^42^. Each sample was mapped against the RSRS downloaded from phylotree.org and checked against haplogroup-defining sites for the sample-specific haplogroup (Data S1A).

Samtools 1.9^39^ option *stats* was used to determine the number of final reads, average read length, average coverage etc. The final average endogenous DNA content of all individuals (proportion of reads mapping to the human genome) was 10.39% (0.09 - 38.82%) (Data S1A).

#### Calculating genetic sex estimation

Genetic sex was calculated using the methods described in^43^, estimating the fraction of reads mapping to Y chromosome out of all reads mapping to either X or Y chromosome. Additionally, sex was determined using a method described in^44^, calculating the X and Y ratio by the division of the coverage by the autosomal coverage. Here, the sex was calculated for samples with a coverage >0.01× and only reads with a mapping quality >10 were counted for the autosomal, X, and Y chromosome (Data S1A).

#### Determining mtDNA haplogroups

Mitochondrial DNA haplogroups were determined using Haplogrep2 on the command line. For the determination, the reads were re-aligned to the reference sequence RSRS and the parameter - rsrs were given to estimate the haplogroups using Haplogrep2^45, 46^(Data S1A). Subsequently, the identical results between the individuals were checked visually by aligning mapped reads to the reference sequence using samtools 0.1.19^39^ command *tview* and confirming the haplogroup assignment in PhyloTree. Additionally, private mutations were noted for further kinship analysis. The polymorphisms were estimated using the online platform of haplogrep2. Here, the variant calling files (vcf) were uploaded to the online platform and the known polymorphism in the RSRS were converted to rCRS (Data S1D).

#### Y chromosome variant calling and haplotyping

A total of 161,140 binary Y chromosome SNPs that have been detected as polymorphic in previous high coverage whole Y chromosome sequencing studies^47–49^ were called in 9 male individuals with more than 0.01× autosomal coverage using ANGSD-0.916^47–49^ ‘-doHaploCall’ option. A subset of 826 sites had at least one of the 9 individuals with a derived allele (DATA C-Table Y). Basal haplogroup affiliations of each sample were determined by assessing the proportion of derived allele calls in a set of primary haplogroup defining internal branches, as defined in^47^, using 1677 informative sites.

#### Kinship analysis

All newly generated individuals with an average human coverage more than 0.03× were selected to assess kinship relationships up to the 3rd degree. We divided the individuals in different groups regarding their geographically area and resulting relationships. First, we were called all individuals using ANGSD-0.916^50^ command --doHaploCall to sample a random base for the positions that are present at MAF>0.1 in the 1000 Genomes GBR population^51^ giving a total of 4,045,514 SNPs for autosomal kinship analysis. The ANGSD output files were converted to .tped format and used as an input for kinship analyses with READ^17^. First, we tested all individuals together and found one 1st degree relationship between ORD001 and ORD009 and two 2nd degree relationships between ORD004, SAL007, and SAL011. We tested those relationships using all individuals from Ordona including SAL007 and SAL011 and all individuals from Salapia including ORD004. We compared the estimated mtDNA haplogroups, radiocarbon dates, genetic sexes, average human coverages, and the age of death from the individuals to confirm or dismiss relationships. (Data S1, Figure S1). Based on the outputs, we dismissed the possible 2nd degree relationships.

#### Dataset preparation and pre-processing for autosomal analysis

We assembled a genome-wide dataset of ancient and modern samples by converting the following datasets, where needed, in PLINK format using *convertf* from the EIGENSOFT 8.0.0^52^ and merging them together with PLINK 1.9^53^: i) the ancient individuals and the Mbuti Congolese individuals (HGDP^54^) from the “1240K” dataset from Dr David Reich laboratory (https://reich.hms.harvard.edu/allen-ancient-dna-resource-aadr-downloadable-genotypes-present-day-and-ancient-dna-data, version 44.3 release), ii) the modern samples from “1240K+HO” of the same release, iii) the Italian Chalcolithic and Bronze Age samples produced by a recent study^11^, iv) the genotypes of an Italian hunter-gatherer from the Late Epigravettian site of Riparo Tagliente dated around 17kya^55^, v) additional genome-wide data of 129 modern-day Italian samples covering the entire country^56^, vi) genome-wide data of 217 present-day Apulian individuals (available at this link https://www.ncbi.nlm.nih.gov/geo/query/acc.cgi?acc=GSE44974) and 48 haploid genomes representing the Eurasian component of modern Ethiopians (here called Amhara_NAF for “Non African Fraction”^24^). We excluded the ancient samples showing issues in their group assignment (i.e., “*Ignore*” in the “1240K” dataset) or in their DNA quality (e.g, those not containing the string “PASS”, with the exception of the *Iceman* individual, or those with less than 5,000 SNPs) and one of a pair of related samples. We further refined the list of ancient samples by retaining only those coming from geographical regions relevant for this study (Data S2)^3, 4, 6, 12, 14–16, 18–23, 25, 42, 44, 57–102^. The same geographical filter was applied on the modern samples from “1240K+HO” and only those flagged as “*PASS (genotyping)*” have been selected in order to minimize variation due to different genotyping techniques (Data S3)^23, 54, 72, 103, 104^. In case of duplicated samples, we selected the item showing the highest number of SNPs among the 1240K SNP set. Finally we added the newly generated ancient Apulian samples to the merge.

We used two different SNPs set for the following analyses. The autosomal “1240K” SNP set (1,150,639 SNPs) was used for the analyses involving just ancient samples, such as the *f*-statistics and qpAdm analyses (see below). Conversely, the autosomal “HO” SNP set was used for those analyses relying also on modern populations (e.g., PCA and ADMIXTURE). In the latter case, we further refine the “HO” SNP set by removing monomorphic SNPs and those with more than 5% missing count in modern populations with PLINK, thus obtaining 503,062 SNPs.

#### Principal component analysis

We performed principal component analysis using the program smartpca implemented in EIGENSOFT software 8.0.0^52^ with the parameters “*lsqproject*” and “*shrinkmode*”. In particular, we projected the ancient samples’ genetic variation onto the principal components inferred from the modern samples of the “1240K+HO” dataset. Modern Italian samples coming from^56^, the newly genotyped ones from modern Apulia and the Amhara_NAF were also projected.

#### Admixture

We used the same dataset of modern and ancient individuals presented in the PCA to explore the genetic components of each individual. For the analysis, we exploited the model-based algorithm implemented in ADMIXTURE^105^ projecting ancient individuals (-P flag) into the genetic structure calculated on the modern dataset, due to missing data in the ancient individuals. We performed unsupervised Admixture for K ∈ {2..10} for modern individuals and used the “per-cluster” inferred allele frequencies to project the ancient individuals. We visualised the Q output using R 3.6^106^.

#### f4 statistics

We performed a series of *f*4 statistics using the program *qpDstat* (option f4mode) implemented in the software ADMIXTOOLS 7.0.2^54^ in different forms. In particular, we grouped together ancient published samples from the same cultural/archeological assemblages (labels used for f3 and f4 analyses can be found in the column “Analysis.Label” of Data S2). Conversely, ancient Apulian samples were both analysed as individual samples and grouped together according to their archeological age (Iron Age samples were grouped together under the acronym “IAA”, Iron Age Apulians). 10 individuals from the Mbuti population from Congo coming from the “1240K” dataset were used as an outgroup.

#### Outgroup f3 statistics

To investigate genetic affinities between ancient Apulian samples with other ancient and modern human populations, we performed a series of Outgroup *f*3 analyses in the form *f*3(X, Y; outgroup=Mbuti) using the qp3Pop function implemented in ADMIXTOOLS 7.0.2^54^. As above-described we used the “Analysis.Label” groups for the ancient samples while we used the “Group.Label” already present in the “1240K+HO” annotation file for modern samples.

We also computed the distance matrices from the Outgroup *f*3 results comparing ancient Apulian samples (both Iron and Middle Age samples) with other published ancient samples by subtracting the f3 values from 1 and we performed a classical (metric) Multi-Dimensional Scaling (MDS) with default options using the function *cmdscale* in the stats package of R (version 4.0.4^107^).

#### qpAdm

We exploited the qpWave/qpAdm framework within the software ADMIXTOOLS 7.0.2 to investigate the genetic composition of ancient Apulian samples and other ancient samples of interest with respect to human ancestries that contributed to coeval European population in the Mediterranean area. In particular, we tested a number of left sources from 2 to 5 in combination with a list of reference sources (right). We used the options ‘‘allsnps=YES’’ and “inbreed=NO” (the last option was used because some populations were composed of just one haploid individual). We tested four different sets of left sources. The “base” set included Western hunter-gatherers (*Loschbour_published.DG*), Anatolian Neolithic (26 samples flagged as “Anatolia_N” in the Analysis.Label column of Data S2), Iranian Neolithic (samples Iran_GanjDareh_N), Caucasus hunter-gatherers (KK1.SG and SATP.SG) and Yamnaya (10 Russia_Samara_EBA_Yamnaya samples). To this set of base sources we separately added as left sources: Minoans (both Greece_Minoan_Lassithi and Greece_Minoan_Odigitria, but taken individually), Amhara_NAF and Roman Republican samples (Italy_Central_Republic_IA). In each run, we consistently used the same set of right sources (if present in the left set, they were removed from the right): Russia_Ust_Ishim_HG_published.DG (which was always kept as the first of the list), Russia_Kostenki14, Russia_AfontovaGora3, Eastern hunter-gatherers (I0061=Russia_HG_Karelia, I0124=Russia_HG_Samara, I0211=Russia_HG_Karelia and UzOO77=Russia_EHG), Spain_ElMiron, Belgium_UP_GoyetQ116_1_published_all, Levant_N (Israel_PPNB), Russia_MA1_HG.SG, Israel_Natufian_published, Czech_Vestonice16 and Caucasus hunter-gatherers.

**Additional resources**

## Acknowledgments

The work was supported by the Estonian Research Council grant PUT (PRG243) (A.S., M.M., C.L.S); the European Union through the European Regional Development Fund (Project No. 2014-2020.4.01.16-0030) (C.L.S., M.M.); the European Regional Development Fund (Project No. 2014-2020.4.01.15-0012) (M.M.) and my UniPd PRID 2019 (S.A., L.P.); the authors would like to thank Dr. Francesco Bertolini for facilitating the research of A.R. in the last stage of the manuscript preparation.

## Author contributions

Conceptualization = L.P., C.L.S.

Formal Analysis = S.A., T.S.

Funding acquisition = M.M., L.P.

Investigation = T.S. (aDNA data generation), A.S. (quality control)

Project administration = C.L.S.

Resources = C.S., N.C., L.M.

Supervision = L.P. and C.L.S.

Visualization = S.A., T.S.

Writing – original draft = S.A., T.S., L.P.

Writing – review & editing = All authors

## Supplementary Data

**Data S1. Sample information (Related to Table 1).** (A) Summary archaeological, mapping, uniparental marker, genetic sex estimation, and damage statistics for samples sequenced in this study. (B) Results of radiocarbon dating of newly generated samples in this study. (C) Binary Y chromosome haplogroup-informative variants found in derived state in at least one of the Iron Age Italian genomes. (D) Mitochondrial variants present in the newly generated sample in this study. Polymorphisms are estimated with haplogrep2. Here, the variant calling file (vcf) is uploaded on the online platform and the input file with the known polymorphism in chrRSRS is converted to rCRS. (E) Comparison of the mean P0 values when running READ on different groupings of all individuals. (F) READ results for individuals when running all individuals with coverage over 0.03x together. (G) READ results for individuals when running all Ordona individuals including SAL007 and SAL011 with coverage over 0.03X together. (H) READ results for individuals when running all Salapia individuals including ORD004 with coverage over 0.03X together.

**Data S2.** List and available information of published ancient samples used in this work.

**Data S3.** List and available information of published modern samples used in this work.

**Data S4.** f4 investigating the genetic relationships among Iron Age Apulians (IAA) and modern Apulians. 14 Apulian samples from Raveane et al. 2019 (Modern_Apulia_GP) were used as Modern Apulians (Data S3).

**Data S5.** qpAdm results with 2, 3 and 4 possible sources for each test model (see STAR methods).

**Data S6.** f4 investigating the genetic relationships among Iron Age Apulians (IAA) and the two Middle Age Apulian samples, the putative Daunians’population of origin (ancient populations coming from Crete, Peloponnese, Croatia and Iron Age Italians) and other ancient human groups. f4 analyses were computed in the form f4(IAA, X; Y, Mbuti) where X is a putative Daunians source and Y is another ancient population.

## Supplementary Materials

**Figure S1.**
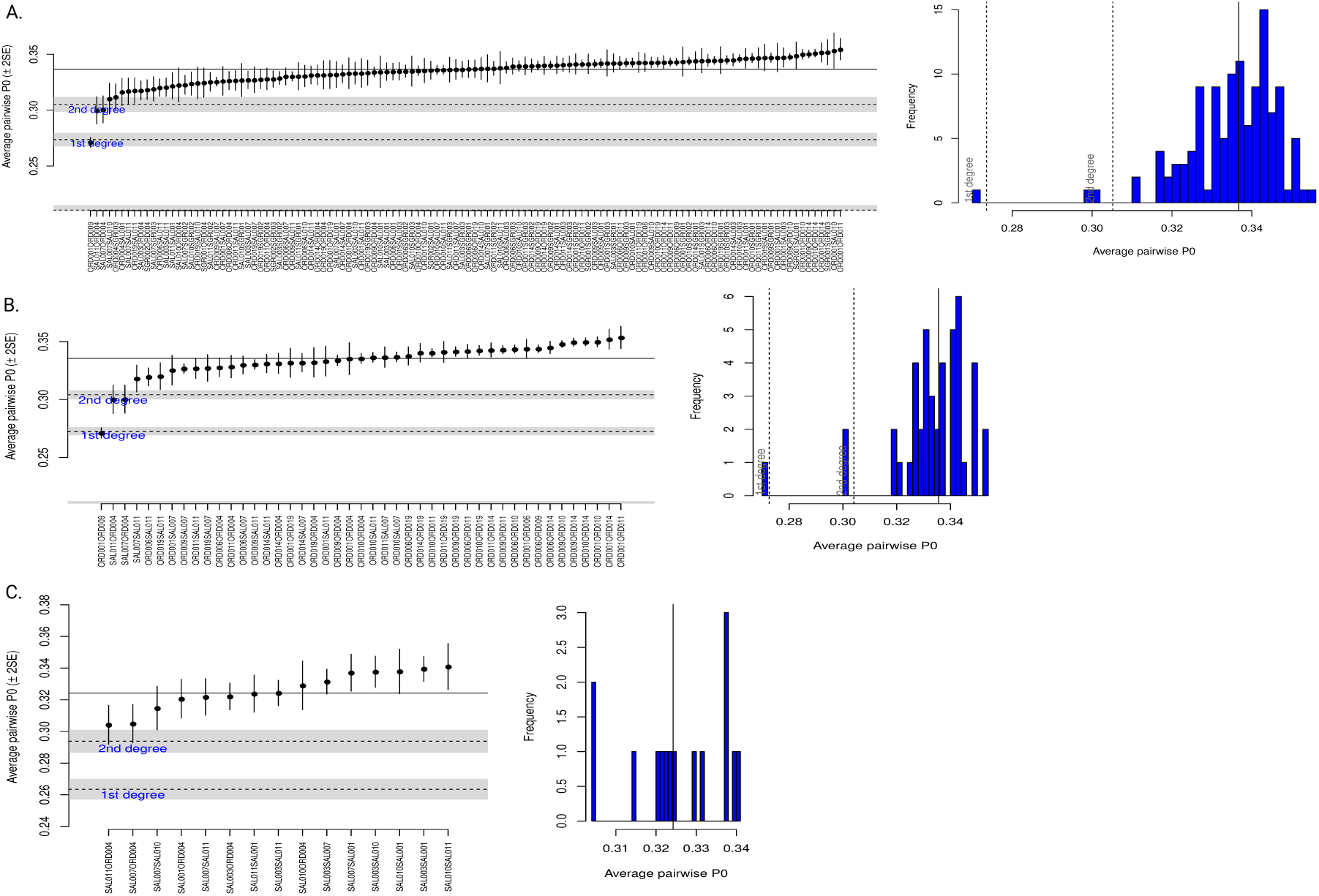
Graphical output of READ. A) All newly generated individuals run together regardless of geography or chronology. B) All newly generated individuals from the archaeological site Ordona plus SAL007 and SAL011. C) All newly generated individuals from the archaeological site Salapia plus ORD004.

**Figure S2.**
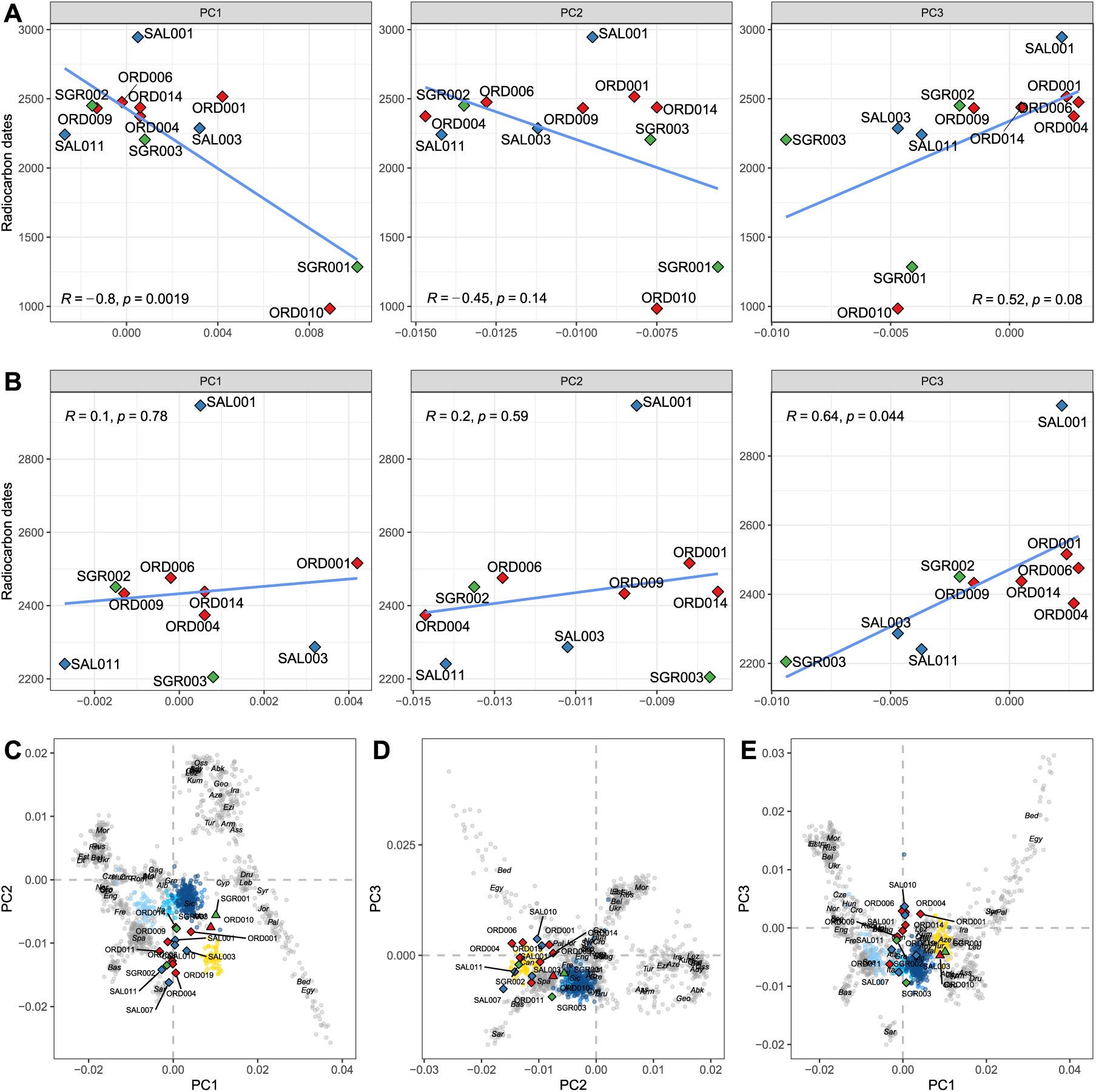
Correlation of radiocarbon dating and principal components for the ancient Apulian samples. A) Pearson correlation of dates with the first three principal components for the radiocarbon dated ancient Apulian samples. B) Same as A, after removing the Middle Age samples ORD010 and SGR001. C), D) and E) show three principal components analyses based on PC1 vs PC2, PC2 vs PC3 and PC1 vs PC3, respectively.

**Figure S3.**
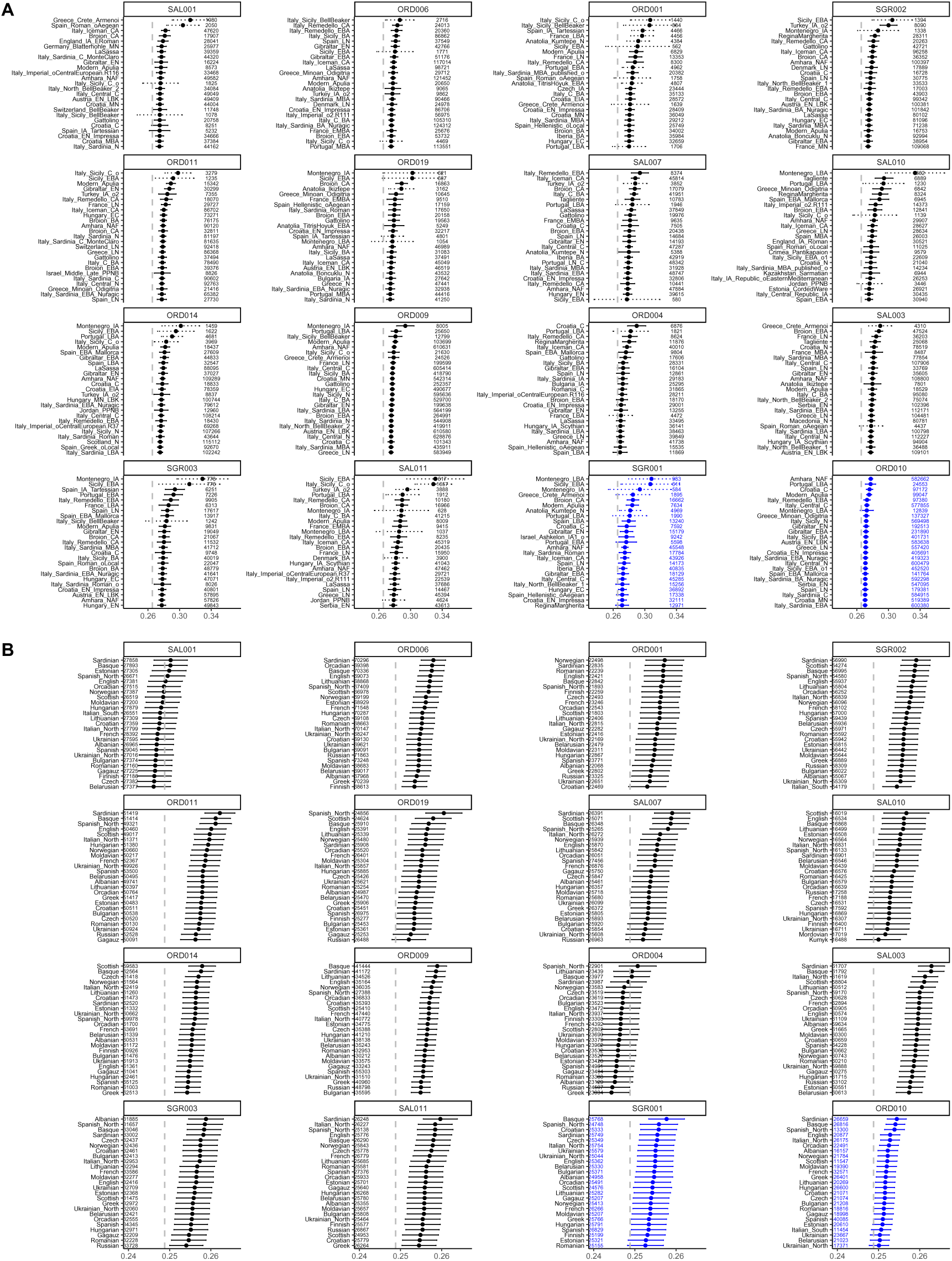
Outgroup *f* 3 analyses in the form *f* 3(Ancient Apulian samples, X; Mbuti). A) Outgroup *f* 3 analyses comparing the newly generated ancient Apulian samples with other ancient human groups (see STAR Methods and Data S1). B) Outgroup *f* 3 analyses comparing the newly generated ancient Apulian samples with present-day populations (see STAR Methods and Data S2). The number of the SNPs used in each run is reported near each *f* 3 value. The ancient Apulian samples are plotted according to their age (SAL001 is the oldest) and Middle Age samples (ORD010 and SGR001) are reported in blue.

**Figure S4.**
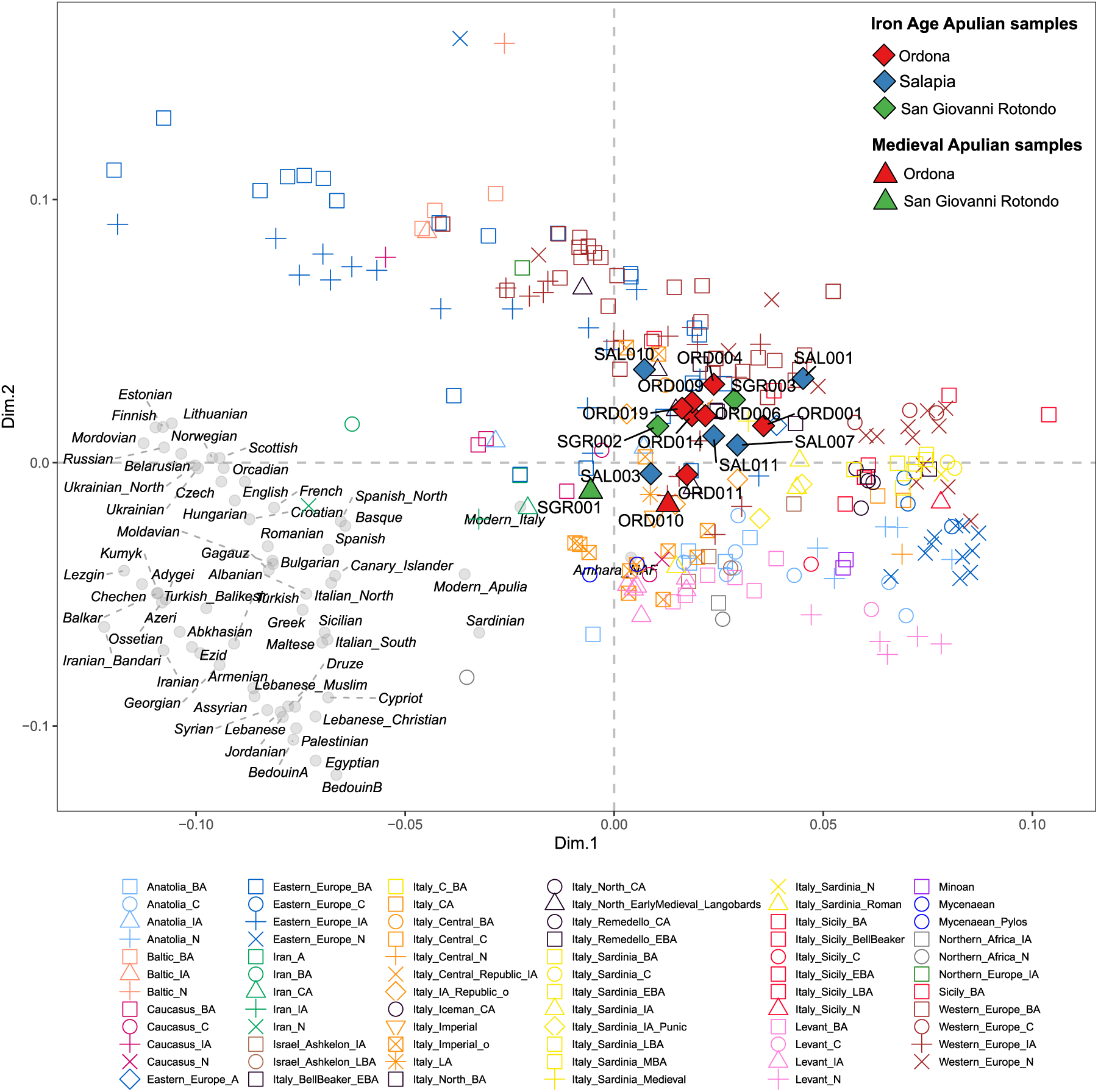
outgroup-*f* 3 multidimensional scaling (MDS) of ancient Apulian samples and published ancient and modern reference populations. A pairwise matrix of outgroup *f* 3(Ancient Apulian samples, X; Mbuti) distance was used to compute the MDS (STAR Methods) after removing samples belonging to Palaelolithic and Mesolithic periods. Ancient and modern groups were aggregated according to their published label (Data S1, Data S2 and STAR Methods).

**Figure S5.**
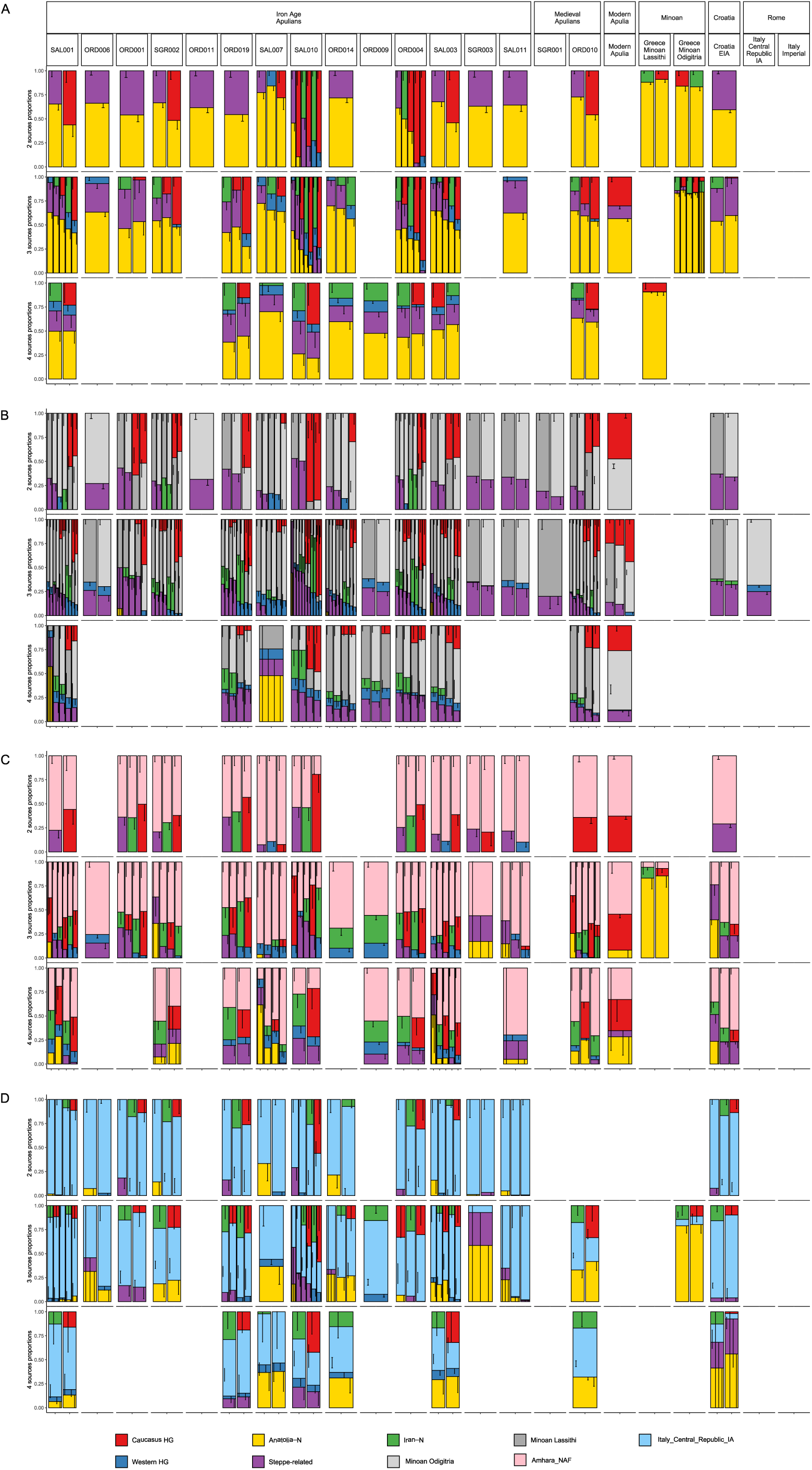
Ancestral composition of ancient Apulian samples inferred by qpAdm. A) qpAdm proportions of ancient Apulian samples and other reference populations (Modern Apulians, Minoans, Croatia EIA and the Roman individuals from the Republican and the Imperial period) using *base* sources: Caucasus and Western hunter-gatherer component (HG), Anatolia Neolithic (Anatolia N), Iranian Neolithic (Iran N), Stepperelated ancestry. B), C) and D) Same as A, using also the Minoans, Amhara NAF and Rome Republicans as source (STAR methods). Only the models with p-values higher than 0.05 are shown.

**Figure S6.**
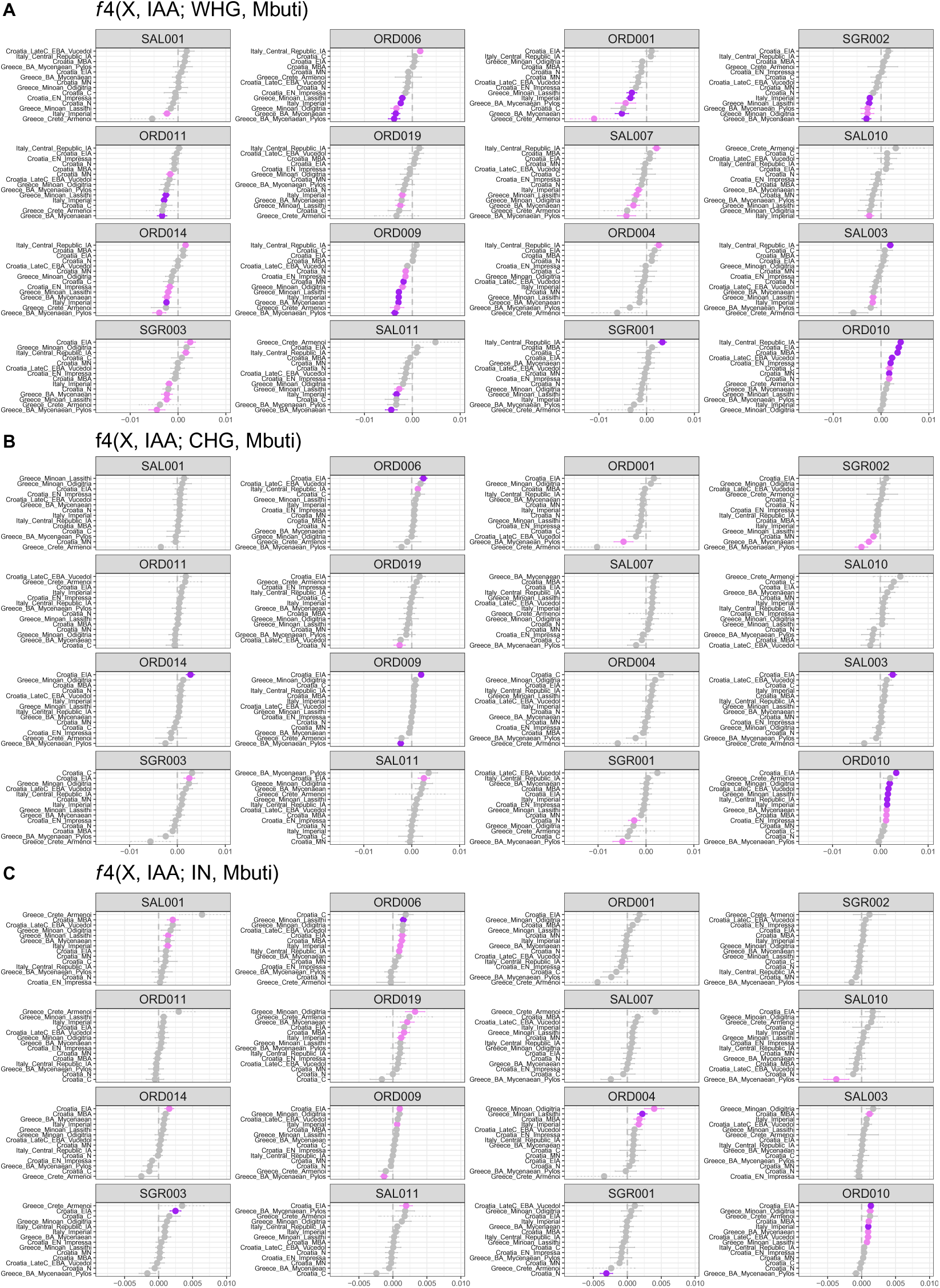
Genetic relationships among Iroan Age Apulians (IAA), their putative sources (ancient populations coming from Crete, Peloponnese, Croatia and Iron Age Italians) and Caucasus, Western hunter-gatherer (CHG and WHG, respectively) and Iranian Neolithic (IN) components. A) *f* 4(X, IAA; WHG, Mbuti) values, where X is a putative source of the Daunian population. B) and C) Same as A, using CHG and IN, respectively. Purple points represent *f* 4 values with significant Z-scores (*Z* 3), pink points represent almost significant *f* 4 values (2 *Z <* 3), while dashed lines indicate *f* 4 runs involving less than 5,000 SNPs. Also the Middle Age Apulian samples (ORD010 and SGR001) were added for comparison.

**Figure S7.**
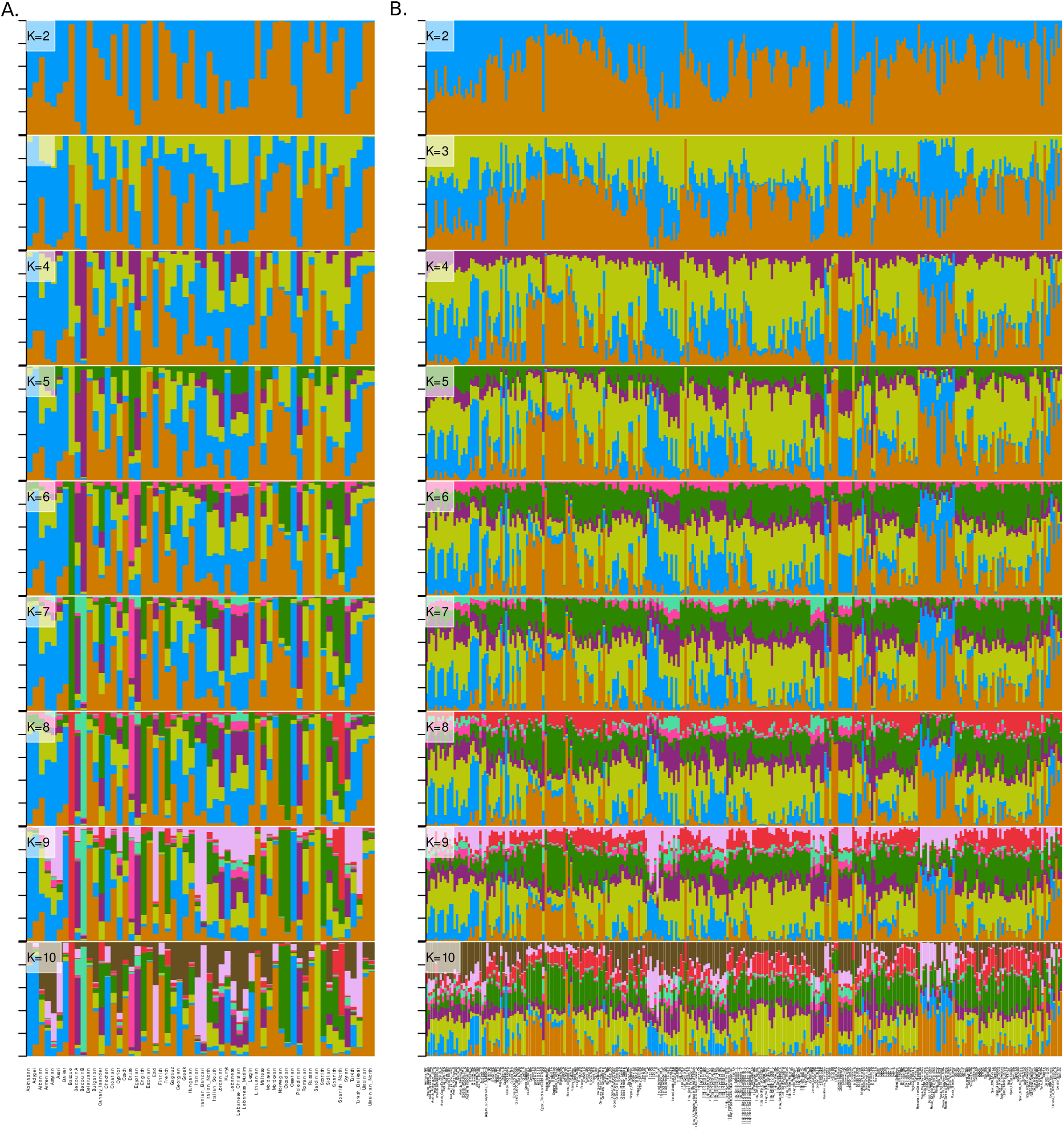
Model-based clustering on modern and ancient individuals as inferred by ADMIXTURE. A) The ancestral components were inferred in a set of 90 modern Eurasian populations. B) Using the allele frequency for ancestral components inferred in A. we projected a subset of ancient Europasian individuals.

**Figure S8.**
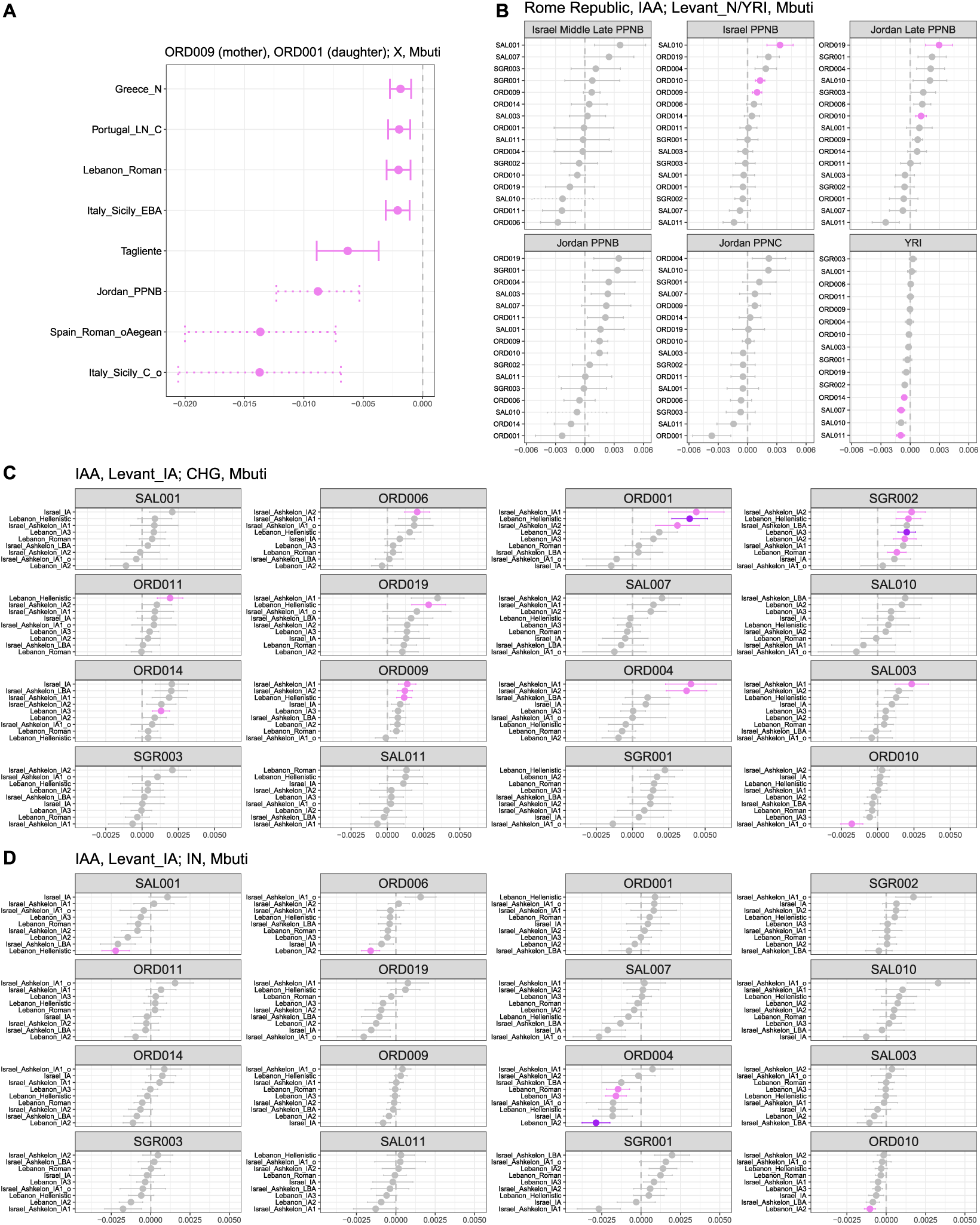
*f* 4 analyses investigating the genetic heterogeneity of IAA samples. A) Almost significant *f* 4 values involving a putative first-degree relationship between ORD009 and ORD001 (possibly, mother and daughter). B) *f* 4 analyses involving Rome Republicans, IAA and Levant N/YRI. C) and D) show the *f* 4 analyses involving IAA, Levant N and, alternatively, CHG and IN. In all plots, purple points represent *f* 4 values with significant Z-scores (*Z* 3), pink points represent almost significant *f* 4 values (2 *Z <* 3), while dashed lines indicate *f* 4 runs involving less than 5,000 SNPs. Also the Middle Age Apulian samples (ORD010 and SGR001) were added for comparison.

